# Rapid processing of observed touch through social perceptual brain regions: an EEG-fMRI fusion study

**DOI:** 10.1101/2023.05.11.540376

**Authors:** Haemy Lee Masson, Leyla Isik

## Abstract

Seeing social touch triggers a strong social-affective response that involves multiple brain networks, including visual, social perceptual, and somatosensory systems. Previous studies have identified the specific functional role of each system, but little is known about the speed and directionality of the information flow. Is this information extracted via the social perceptual system or from simulation from somatosensory cortex? To address this, we examined the spatiotemporal neural processing of observed touch. Twenty participants watched 500 ms video clips showing social and non-social touch during EEG recording. Visual and social-affective features were rapidly extracted in the brain, beginning at 90 and 150 ms after video onset, respectively. Combining the EEG data with fMRI data from our prior study with the same stimuli reveals that neural information first arises in early visual cortex (EVC), then in the temporoparietal junction and posterior superior temporal sulcus (TPJ/pSTS), and finally in the somatosensory cortex. EVC and TPJ/pSTS uniquely explain EEG neural patterns, while somatosensory cortex does not contribute to EEG patterns alone, suggesting that social-affective information may flow from TPJ/pSTS to somatosensory cortex. Together, these findings show that social touch is processed quickly, within the timeframe of feedforward visual processes, and that the social-affective meaning of touch is first extracted by a social perceptual pathway. Such rapid processing of social touch may be vital to its effective use during social interaction.

**Significance Statement:** Seeing physical contact between people evokes a strong social-emotional response. Previous research has identified the brain systems responsible for this response, but little is known about how quickly and in what direction the information flows. We demonstrated that the brain processes the social-emotional meaning of observed touch quickly, starting as early as 150 milliseconds after the stimulus onset. By combining EEG data with fMRI data, we show for the first time that the social-affective meaning of touch is first extracted by a social perceptual pathway and followed by the later involvement of somatosensory simulation. This rapid processing of touch through the social perceptual route may play a pivotal role in effective usage of touch in social communication and interaction.

## Introduction

Touch evokes a strong social and emotional response in third-party observers as well as the direct recipient (Hertenstein et al., 2006a, 2009). During mere observation, humans accurately extract the social-affective meaning of a touch gesture, with high inter-observer reliability (Lee Masson and Op de Beeck, 2018). The functional magnetic resonance imaging (fMRI) literature suggests viewing social touch increases posterior insula responses to observed social touch (Morrison et al., 2011, but see Ebisch et al., 2011) and leads to shared somatosensory responses between self-experienced and observed social touch (Ebisch et al., 2008, 2011; Gazzola et al., 2012). Further, the social-affective meaning of observed touch is represented in social-cognitive brain areas, including temporoparietal junction and posterior superior temporal sulcus (TPJ/pSTS), as well as somatosensory cortex, and observing social touch leads to enhanced functional communication between these regions (Lee Masson et al., 2018, 2020).

These results suggest that observed touch is understood not only from direct perceptual signals, but also via somatosensory simulation (see the review by Peled-Avron and Woolley, 2022). Somatosensory simulation has been shown to vary greatly between individuals based on the degree of emotional empathy, attitude towards social touch, and autistic traits (Gallese and Ebisch, 2013; Giummarra et al., 2015; Peled-Avron et al., 2016; Peled-Avron and Shamay-Tsoory, 2017; Lee Masson et al., 2018, 2019). However, recent work has called into question the direct role of simulation in other aspects of social perception like action recognition (Caramazza et al., 2014). Due to the slow temporal resolution of fMRI, it is difficult to understand the direction of information flow between somatosensory and social perceptual brain regions.

Prior electroencephalogram (EEG) studies have investigated the neural processing of social touch observation, mostly focusing on the mu rhythm indexing somatosensory simulation and event-related potentials (ERPs) (Peled-Avron et al., 2016; Schirmer and McGlone, 2018; Addabbo et al., 2020). A few prior studies have provided initial insight into how fast the brain processes an observed touch event, and found the observation of another person receiving simple, non-social touch, such as a paintbrush touching a hand, evoked early involvement of somatosensory ERPs (Adler and Gillmeister, 2019; Rigato et al., 2019b, 2019a). In contrast, adding social-affective complexity to a touch scene results in longer processing time reflected by increases in P100 and late positive potential (Peled-Avron and Shamay-Tsoory, 2017; Schirmer and McGlone, 2018). However, no prior study on social touch has directly linked stimulus features to EEG timeseries, so it remains to be seen how quickly visual and social-affective features of observed touch are processed. Furthermore, although the involvement of the somatosensory cortex has been suggested in these EEG studies, spatial localization with EEG is often inconclusive.

Here we ask whether social touch features are processed via social perception or somatosensory simulation. To answer this question, we apply new methods in fMRI-EEG fusion (Cichy and Oliva, 2020), and use representational similarity analysis (RSA) to link stimulus features, fMRI multi-voxel patterns, and EEG activity patterns from observed touch scenes. In particular, we examine how fast each social-affective feature is processed, as well as the time course of feature representations in different brain regions, including early visual cortex (EVC), TPJ/pSTS and somatosensory cortex. If somatosensory simulation drives social touch perception, we would expect the neural patterns from somatosensory cortex to correlate with earlier EEG activity patterns than TPJ/pSTS. However, we find the opposite results: EEG signals correlate first with EVC followed by TPJ/pSTS, and finally somatosensory cortex. We further find that while EVC and TPJ/pSTS each share unique variance with EEG neural patterns and social-affective features, somatosensory cortex does not. Together, these results indicate that social-affective features in observed touch scenes are directly extracted via a social perceptual pathway without direct somatosensory simulation.

## Materials and Methods

### Participants

21 participants (male = 7, mean age = 20.9 years, age range 18 - 32 years) took part in the EEG study. They all reported normal or corrected-to-normal vision. 20 participants were recruited through the Johns Hopkins University SONA psychological research portal and received research credits as compensation for their time. One participant was compensated with a monetary reimbursement. They provided written informed consent before the experiment. The study was approved by Johns Hopkins University Institutional Review Board (protocol number HIRB00009835).

### Stimuli

We used a stimulus set developed and validated in a previous study (Lee Masson and Op de Beeck, 2018). The original stimulus set consisted of 39 social and 36 non-social three-second videos. Social videos showed 18 pleasant, 3 neutral, and 18 unpleasant human-human touch interactions, such as hugging or slapping a person. Non-social videos showed human-object touch manipulation that are matched to the social videos in terms of biological motion, such as carrying a box or whacking a rug (Figure 1).

**Figure 1.**
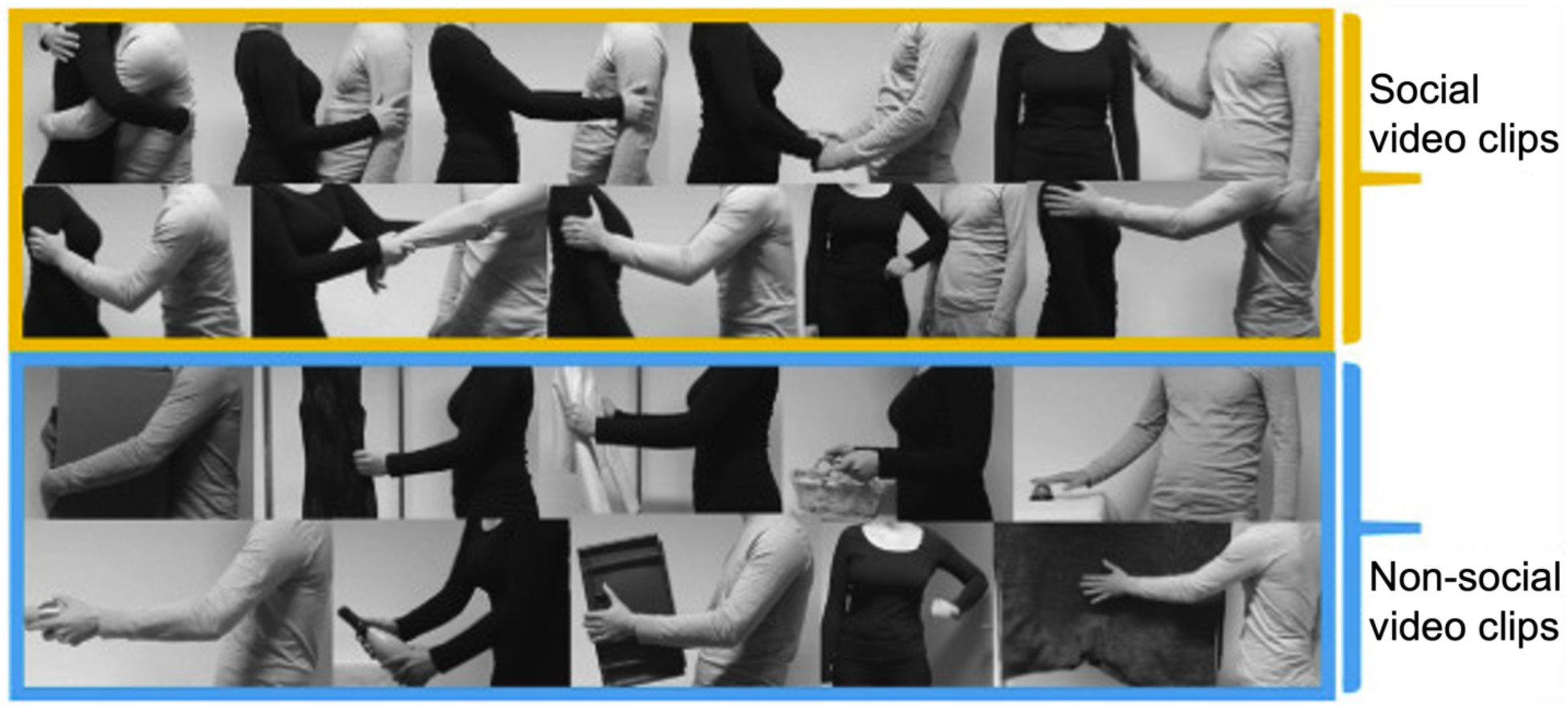
A few example frames of video clips in 2 categories (social and non-social). Images in a yellow box show representative frames of social touch videos. The top row shows examples of pleasant touch events, and the second row unpleasant touch events. Frames of non-social touch videos displaying matched human-object interactions are shown in a blue box. This figure is published in compliance with a CC-BY-NC-ND license (https://creativecommons.org/licenses/by-nc-nd/4.0/) and is re-used from Fig. 1 in the original study (Lee Masson et al., 2018). The complete set of original video materials is available at https://osf.io/8j74m/.

For the current study, all videos were trimmed to a duration of 0.5 s centered around the touch action to improve time-locking to the EEG signal. Each video contained 13 frames of 720 (height) × 1280 pixels in size. To ensure trimming the videos did not alter the perceived valence and arousal of touch events, we had human annotators from the online platform Amazon Mechanical Turk rate the valence and arousal of each video. We found ratings were highly correlated between the trimmed and full-length videos (Spearman r = 0.96 for valence and 0.94 for arousal).

### EEG experimental procedure and design

During the EEG recording, participants were seated comfortably on a chair and viewed the videos displayed on a back-projector screen in a Faraday chamber (visual angle: 15 × 13 degrees, distance from the screen: 45 cm). They were instructed to view each video and press a button on a Logitech game controller when they detected two consecutive videos that were identical (One-Back task). These catch trials, involving participants’ motor responses, were excluded from the analysis. This orthogonal task aimed to get participants to pay attention to the videos. Each block (N = 15) consisted of 83 trials, with the 75 videos presented once in a pseudo-random order and four catch trials (i.e., four randomly selected videos were shown twice).

The videos were shown using an Epson PowerLite Home Cinema 3000 projector. A light-sensitive photodiode was used to track the onset and offset of video presentation on the projector screen to account for delays between the time that the computer initiated stimulus presentation and the time that the stimulus was displayed. Each trial started with a black fixation cross on a white background screen, shown for a random duration between 1 and 1.5 s, followed by a 0.5 s video. The total duration of each block was approximately 2.4 min. After each block, participants were encouraged to take a break for as long as needed before continuing with the following block. The experiment consisted of 1245 trials and took less than 1 hour, including breaks.

The experiment was run using the Psychophysics Toolbox (Brainard, 1997; Pelli, 1997; Kleiner et al., 2007) in MATLAB (R2020a, The Mathworks, Natick, MA). The EEG experiment employed a within-subject design, with EEG neural activity as the dependent variable, and stimulus features and fMRI neural patterns as the independent variables.

### EEG acquisition and preprocessing

During the experiment described above, the EEG data were continuously collected with a sample rate of 1000 Hz using a 64-channel Brain Products ActiCHamp system with actiCAP electrode caps (Brain Products GmbH, Gilching, Germany). An electrolyte gel was applied to each electrode to improve impedances. We aimed to keep electrode impedances below 25 kΩ throughout the experiment. The Cz electrode acted as an online reference.

MATLAB R2020a and the FieldTrip toolbox were used for EEG data preprocessing (Oostenveld et al., 2011). First, we corrected for lags between the stimulus triggers and the stimulus presentation on the projector screen by aligning the EEG data to the stimulus onset defined by the photodiode. The data were segmented into 1.2 s epochs (0.2 s pre-stimulus to 1 s post-stimulus onset). Next, the data were baseline-corrected using the 0.2 s pre-stimulus and high pass filtered at 0.1 Hz to remove slow drifts.

For artifact rejection, we discarded bad channels and trials contaminated with muscle or eye artifacts. Data were band-pass filtered from 110 to 140 Hz, and a Hilbert transformation was applied. Timepoints with a z-value above 15 were considered to belong to muscle artifacts and removed. In addition, channels and trials with high variance were manually rejected using the ft_rejectvisual function in FieldTrip. Afterward, independent component analysis was performed to detect eye movement components and remove eye artifacts from the data. Catch trials and any trials with participants’ motor responses were excluded from the analysis. This preprocessing step yielded, on average, 1122 ± 50 trials and 62.6 ± 1.27 channels. Only participants with no more than six channels removed were kept, resulting in one participant being excluded from further analysis. Lastly, preprocessed data were re-referenced to the median across all channels, low-pass filtered at 100 Hz, and downsampled to 500 Hz.

### Event-related potential analysis

We performed ERP analysis to investigate when and where the brain shows different neural responses to social versus non-social touch. ERPs from all trials were averaged for each condition and participant using the ft_timelockanalysis function in the FieldTrip toolbox. For group-level analysis, the grand averaged ERPs over participants for each condition were computed using the ft_timelockgrandaverage function. As described above, noisy channels were excluded from each participant’s data, and group-level ERP analysis included 48 channels common to all 20 participants. Differences between the two conditions were calculated and visualized with the ft_math and ft_topoplotER functions, respectively. For statistical inference, ERPs were averaged across successive 100 ms time slices for each channel, from 0.2 s pre-stimulus to 1 s post-stimulus onset. A t-test was performed with the Bonferroni correction at an alpha level of 0.05 to determine brain response differences between the two conditions using the ft_timelockstatistics function, and we report Z-scored T-values.

### Pairwise EEG decoding analysis to construct representational dissimilarity matrices

We performed time-resolved multivariate pattern analysis to link the EEG neural activity patterns to 1) stimulus features and 2) the fMRI multi-voxel patterns of the early visual cortex (EVC), social brain regions: TPJ/pSTS, and the somatosensory cortex (Figure 2).

**Figure 2.**
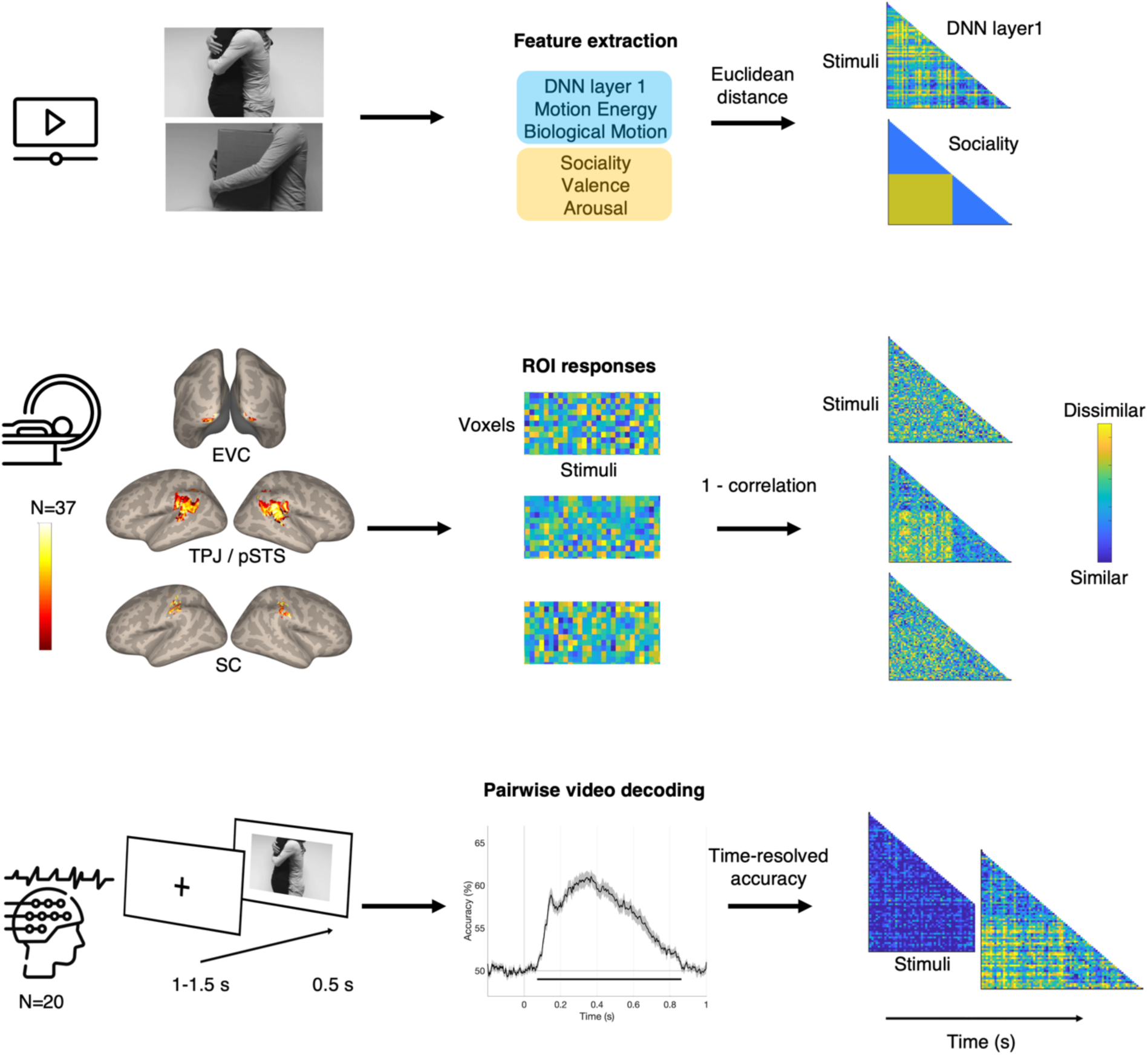
Experimental and analysis overview for evaluating how EEG signals correlate to six stimulus features as well as the fMRI multi-voxel patterns of three ROIs. Top panel: three visual and three social-affective features were extracted from each video. Euclidean distance was calculated between all pairs of stimuli to generate representational distance matrices (RDM) for each feature. DNN layer 1 and sociality RDMs are visualized. Blue in the matrices indicates high similarity while yellow indicates high dissimilarity for each stimuli pair. Middle panel: Activated voxels during video viewing in three ROIs are shown. Voxels in bright yellow in the brain map indicate activations found in all 37 fMRI participants while voxels in dark red indicate activations found in one participant. Across these voxels, dissimilarity (1-correlation) was calculated between all pairs of stimuli to generate fMRI RDMs. RDMs of three ROIs are visualized. Bottom panel: EEG signals from 21 participants were recorded during video viewing. Pairwise decoding accuracy was calculated at each timepoint with a 2 ms resolution and used to generate time-resolved EEG RDMs. The horizontal line in the decoding accuracy plot marks significant time windows where observed accuracy is above chance (one-tail sign permutation for group-level statistics, cluster-corrected p < 0.05). EEG RDMs at 0 ms and 200ms post-stimulus onset are visualized. EVC = early visual cortex, TPJ/pSTS = temporoparietal junction/posterior superior temporal sulcus, SC = somatosensory cortex.

Before decoding, each participant’s preprocessed EEG data were randomly split into two folds for cross-validation. To improve the signal-to-noise ratio (Dima et al., 2022), we created pseudo-trials by averaging 6∼8 trials corresponding to the same stimulus. Multivariate noise normalization was also performed (Guggenmos et al., 2018). Finally, time-resolved pairwise EEG decoding analysis was performed using a linear support vector machine classifier implemented in the LibSVM library (Chang and Lin, 2011). Voltages from all EEG channels were considered features at each time point. The process was repeated ten times, and decoding accuracies were averaged over all iterations for each participant. Decoding performance was used to create a time-resolved EEG neural RDM, in which high decoding accuracy results in patterns that are more different from one another while low accuracy results in patterns that are more similar (Figure 2). Since decoding is only performed to generate the RDM, decoding results are not reported in the results section. Instead, Figure 2 at the bottom panel includes the group-level decoding results, averaged across all pairs of stimuli and participants. Note that the time course of group averaged decoding accuracy was similar to results observed in previous studies using visual stimuli (Isik et al., 2020; Dima et al., 2022).

### EEG to feature representational similarity analysis

Pairwise decoding results were used to generate each participant’s time-resolved neural EEG RDMs, which were correlated to six stimulus features and three brain regions from the fMRI data. Features included three visual features (a low-level visual feature, motion energy, and perceived biological motion similarity between video pairs) and three social-affective features (sociality, valence, and arousal). How each of the six feature RDMs was generated is briefly described below.

1) The low-level visual feature was extracted from the first convolutional layer of AlexNet (Krizhevsky et al., 2012), pretrained on the ImageNet dataset (Russakovsky et al., 2015), using PyTorch (version 1.4.0). Layer 1 was chosen as it captured early visual responses well in a previous EEG study that also used video clips depicting everyday actions (Dima et al., 2022). The middle frame of each 0.5 s video clip was normalized, resized to 640 x 360 pixels, and became an input to the first layer of AlexNet. The size of the kernels for the first layer is 11 x 11, resulting in an output size of 89 x 159 x 64 for each stimulus. The Euclidean distance between the resulting features for each pair of videos was used to generate the low-level visual RDM. 2). For motion energy, we estimated optical flow for each pixel of every video frame using the Farneback method implemented in MATLAB. The sum of optic flow across all pixels for each frame was calculated for each video (13 frames x 75 videos). The Euclidean distance between all video pairs was used to generate a motion energy RDM. 3). In a separate study (unpublished, approved by the Social and Societal Ethics Committee of KU Leuven (G-2016 06 569)), 45 participants who provided written informed consent before the experiment viewed all pairs of videos and made judgments on biological motion similarity using a 7-point Likert scale (“How similar are the movements of human touches in the two videos?” 1 - very distinct, 7 - identical). The pairwise similarity ratings were averaged across participants and subtracted by 7. The resulting scores were used to generate a biological motion RDM.

4) The sociality of each video, here defined as the social versus non-social content of touch, is a binary feature. The distance between pairs of videos from the same category was expressed as 0 (same), while videos from different categories were assigned distances of 1 (different) in the matrix. 5) For valence and 6) arousal of touch, we re-used ratings of 37 participants from our two previous studies (Lee Masson and Op de Beeck, 2018; Lee Masson et al., 2019) and generated both RDMs by calculating the absolute value of the rating differences for each pair of stimuli.

A rank correlational method was used to link each participant’s time-resolved EEG neural RDMs to each stimulus feature. Correlation analysis was performed using 10 ms sliding windows with an overlap of 6 ms of EEG neural activity. We calculated leave-one-subject-out correlation where each subject’s EEG signals were correlated with the group average (excluding that subject) and the average across held out subjects was used as a measure of the noise ceiling. (Nili et al., 2014).

### EEG to fMRI Representational Similarity Analysis

To examine spatio-temporal neural dynamics during touch observation, we correlated time-resolved EEG neural RDMs to fMRI activity RDMs. fMRI data were collected in our previous studies where 37 neurotypical adults viewed the original version of the touch stimuli (3s video clips) and received positive and negative affective touch during a somatosensory localizer scan (Lee Masson et al., 2018, 2019). Full details on fMRI data acquisition and preprocessing procedure can be found in our previous study (Lee Masson et al., 2018). fMRI data analysis related to the current RSA methods is summarized below.

Neural RDMs of three regions of interest (ROI) – EVC, TPJ/pSTS, and somatosensory cortex – were included in the current study. For EVC and TPJ/pSTS, visually responsive voxels (i.e., voxels showing increased responses to videos versus rest) were selected within a corresponding anatomical template, i.e., Brodmann area 17 from the SPM Anatomy toolbox (Eickhoff et al., 2005), and TPJ from connectivity-based parcellation atlas (Mars et al., 2012). We name the latter region TPJ/pSTS as the TPJ template includes voxels that are also part of pSTS, and our voxel definition is likely to extract perceptual voxel responses. To define an ROI involved in somatosensory simulation, a separate touch localizer was used to identify voxels that respond to self-experienced affective touch within an anatomical template, Brodmann area 2 from the SPM Anatomy toolbox. Identified voxels in the somatosensory cortex represent the social-affective meaning of observed touch and are hence involved in somatosensory simulation (Lee Masson et al., 2018, 2019). The pairwise correlation between all voxels in each ROI was used to create fMRI neural RDMs and capture the differences in multi-voxel neural response patterns between each video pair. A rank correlational method was used to link each participant’s time-resolved EEG neural RDMs to each ROI.

Figure 3 shows pairwise correlations between the RDMs of six features and three ROIs from the fMRI data. A strong correlation between sociality and perceived arousal of touch was observed (r = 0.64). As found in the previous study (Lee Masson et al., 2018, 2019), TPJ/pSTS strongly represents sociality of observed touch (r = 0.51), with this region showing significant neural pattern similarity with the somatosensory cortex (r = 0.26).

**Figure 3.**
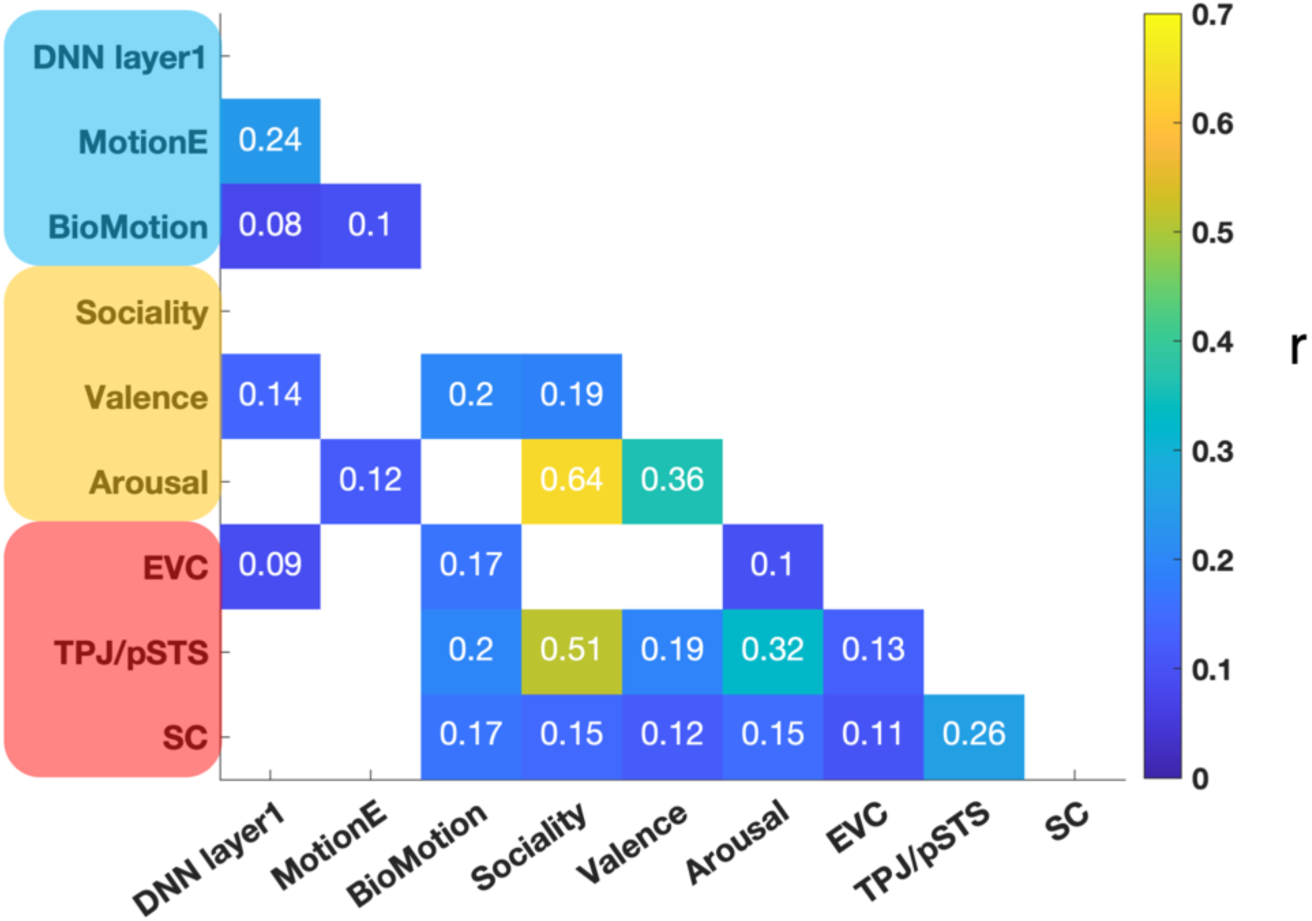
Pairwise correlations between predictors. Three visual (labels colored in blue) and three social-affective (labels colored in orange) features, and fMRI responses of three ROIs (labels colored in red) are included in the RSA. Lower diagonal cells in the matrix contain information about correlation coefficients (*r*) from pairwise comparisons. White colored lower diagonal cells indicate no significant correlation between predictors. No significant negative correlation was observed. DNN = deep neural network, MotionE = motion energy, BioMotion = biological motion, EVC = early visual cortex, TPJ/pSTS = temporoparietal junction/posterior superior temporal sulcus, SC = somatosensory cortex

### Variance partitioning

A time-resolved variance partitioning approach was adopted to examine 1) the unique contribution of the three key brain regions to the EEG signal to characterize the direction of information flow between them, and 2) their shared contribution with the sociality feature to EEG signals. We focus on the sociality feature as TPJ/pSTS and somatosensory cortex both represent the social content of observed touch (Lee Masson et al., 2018, 2019).

To this end, we fit seven different multiple regression models with every possible combination of the three ROIs: EVC, TPJ/pSTS, and somatosensory cortex (i.e., each ROI alone, all three combinations of pairs, and the combination of all three), as well as seven combinations of sociality with the two key ROIs: Sociality, TPJ/pSTS, and somatosensory cortex. The time-resolved EEG neural RDM averaged across subjects was the dependent variable in all models. All variables were normalized prior to regression analysis. Resulting R^2^ values from seven regression models were used to calculate both unique and shared variance. The amount of unique variance explained by each predictor was calculated as follows:

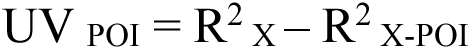

R^2^ is a goodness-of-fit measure for the regression model, representing the amount of variance explained by a model consisting of chosen predictors. X reflects three selected predictors included in the model (e.g., brain regions). POI reflects a predictor of interest (e.g., EVC). X-POI reflects the remaining predictors without the predictor of interest (e.g., TPJ/pSTS, and somatosensory cortex). UV _POI_ is the amount of unique variance explained by POI.

The amount of shared variance (SV) explained by every possible combination of selected predictors was calculated as follows:

_1)_ SV _123_ = R^2^ _12_ – R^2^ _2_ + R^2^ _3_ – R^2^ _12_ – R^2^ _13_ – R^2^ _23_ + R^2^ _123_
_2)_ SV _12_ = R^2^ _13_ – R^2^ _3_ + R^2^ _23_ – R^2^ _123_
_3)_ SV _13_ = R^2^ _12_ – R^2^ _2_ + R^2^ _23_ – R^2^ _123_
_4)_ SV _23_ = R^2^ _12_ – R^2^ _1_ + R^2^ _13_ – R^2^ _123_

Numbers next to SV and R^2^ denote selected predictors. For example, SV_123_ is the amount of shared variance explained by all three selected predictors. R^2^ _12_ is the amount of variance explained by a model consisting of the first and the second predictor.

### Statistical Analysis

All time-resolved RSA results were tested against chance using a one-tailed sign permutation test (5000 iterations). Multiple comparisons across time were controlled by applying cluster correction with the maximum cluster sum across time windows and an alpha level of 0.05.

### Data and Code Accessibility

The current study used analysis code for EEG preprocessing and multivariate analysis used in a previous study (Dima et al., 2022), available on GitHub (https://github.com/dianadima/mot_action/tree/master/analysis/eeg). The code for stimulus presentation and ERP analysis is available on GitHub (https://github.com/haemyleemasson/EEG_experiment). EEG data are available at https://osf.io/5ntcj/.

## Results

### Early evoked response differences between social and non-social touch

We compared the ERPs evoked by observed social and non-social touch events to measure the effect of sociality on the magnitude of ERPs for each channel and time window. We observed widespread differences across the scalp, beginning at 100 ms post video onset. Anterior sensors showed enhanced activation during social touch observation (regions colored in yellow in Figure 4), whereas activations were stronger at the posterior sensors during non-social touch observation (blue in Figure 4), perhaps due to the presence of objects in these videos. In particular, in most of the time windows, observing social touch evoked the strongest activation at channel F2, located approximately near superior frontal gyrus (Scrivener and Reader, 2022), whereas P8, located near lateral occipital cortex, was most activated in non-social touch. These ERP results indicate that the sociality of video clips affects activations even at early stages of processing.

**Figure 4.**
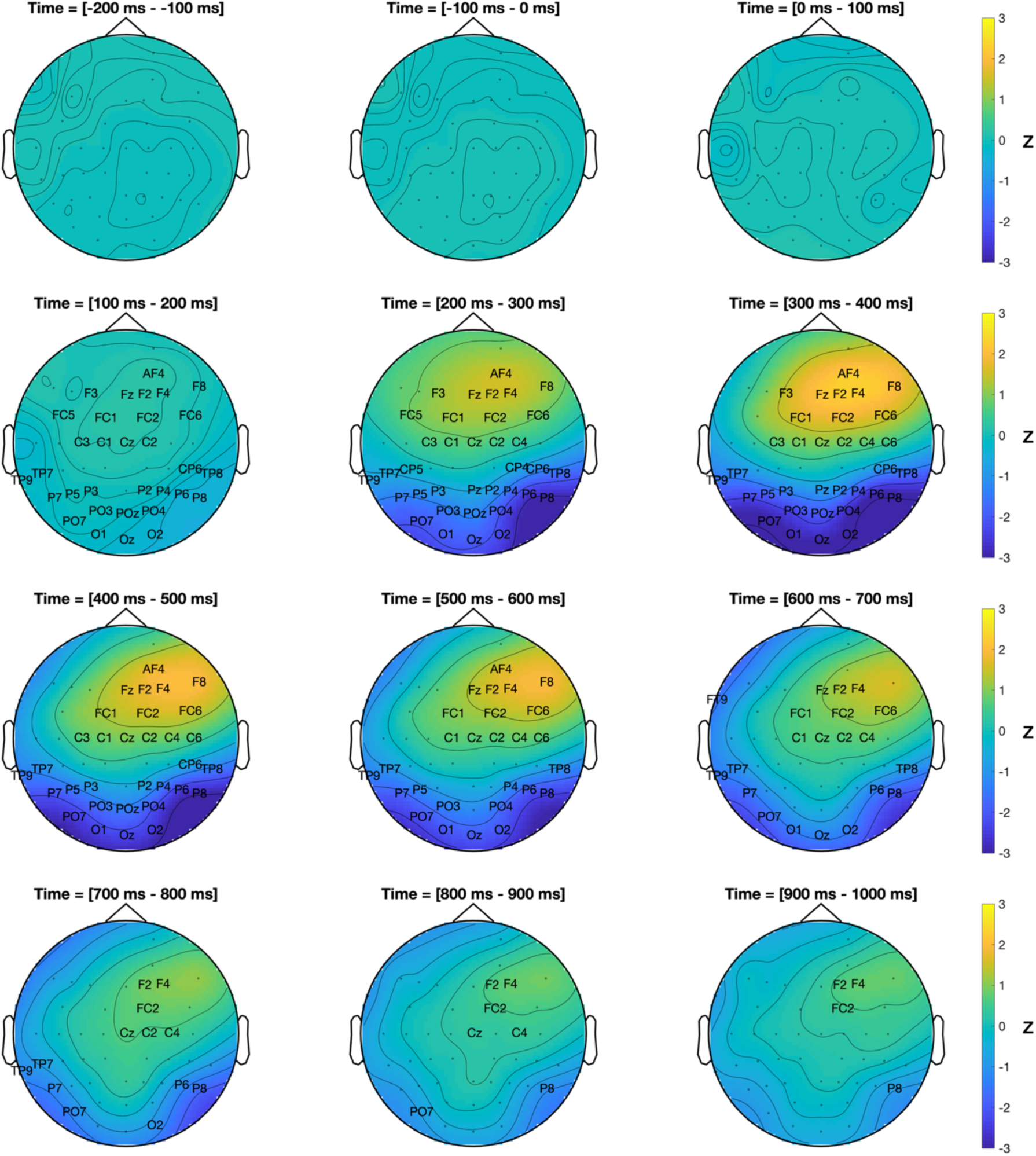
ERP differences between social and non-social touch observation. Topographical scalp maps show the social > non-social contrast, averaged across successive 100 ms time slices. Positive z-scores (increased ERP for social touch) are shown in yellow and negative z-scores (increased ERP for non-social touch) are shown in blue. Channels showing significant differences in ERPs were labeled with channel names in the scalp map. Channels that did not reach statistical significance were marked with dots to indicate their locations on the scalp.

### Social-affective features are processed shortly after visual features

Using time-resolved RSA, we evaluated the neural dynamics of touch observation or how quickly different stimulus features are processed in the brain. To this end, we correlated three visual and three social-affective features to each participant’s EEG neural patterns. A group-level RSA revealed that low-level visual features captured by the first layer of AlexNet correlated significantly with EEG neural patterns beginning at 90 ms post stimulus onset, followed by biological motion and motion energy at 120 and 160 ms, respectively (Figure 5, top). Strikingly, social-affective features correlated with EEG neural patterns shortly after visual features, beginning at 150 ms post video onset for sociality, 170 ms for arousal, and 180 ms for valence (Figure 5, middle). All six features were spontaneously extracted in the brain as participants were not directed to any of these features during the EEG experiment.

**Figure 5.**
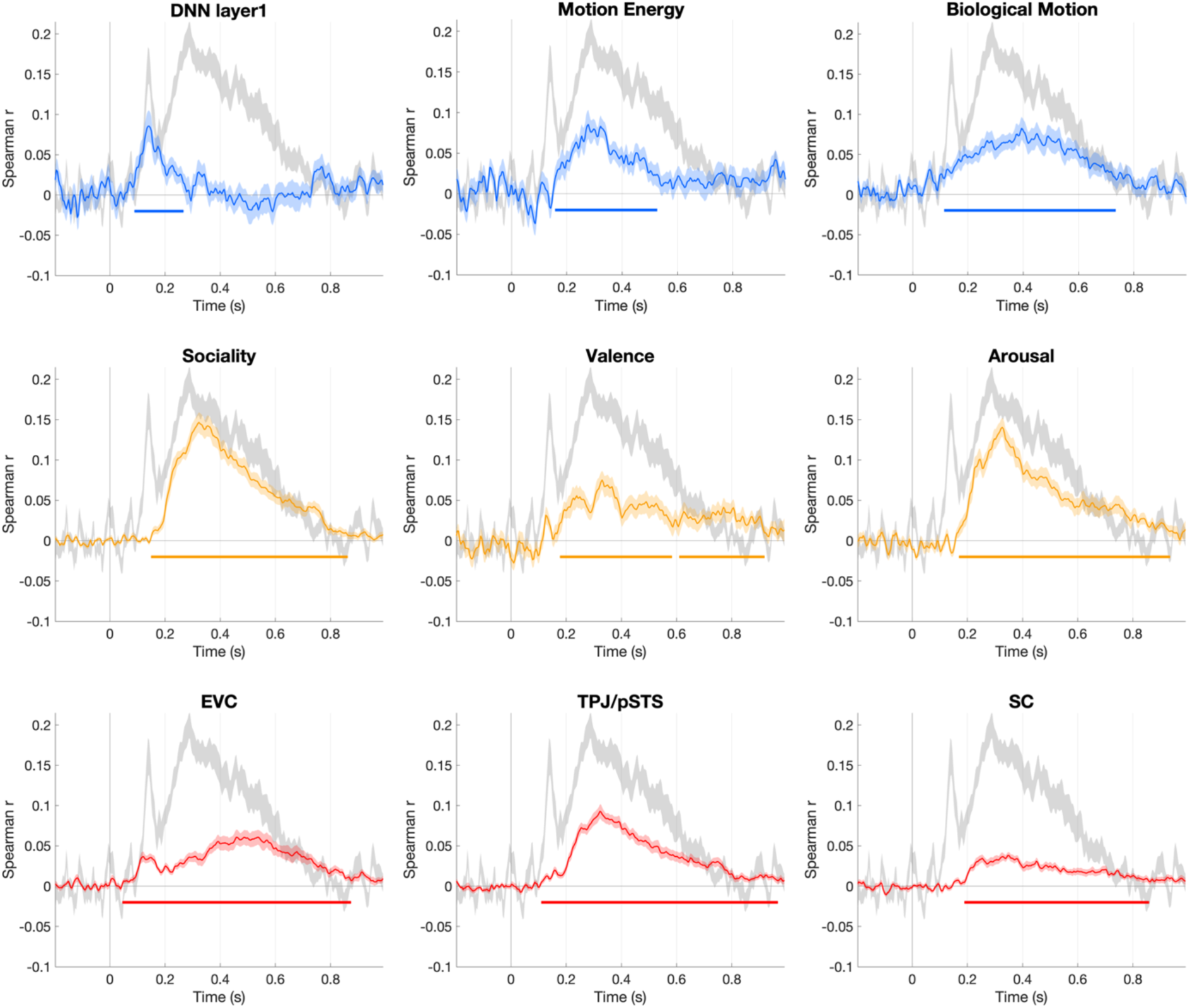
The time courses of all predictor correlations with EEG data (mean ± standard error of the mean (SEM) shown in light colors). This figure shows group-level spearman correlations between visual (time courses shown in blue, top panel), social-affective (time courses in yellow, middle panel), and fMRI activity predictors (time courses in red, bottom panel) and the time-resolved EEG neural patterns. The colored horizontal line in each plot marks significant time windows where observed correlation coefficients are significantly greater than 0 (one-tail sign permutation for the group-level statistics, cluster-corrected p < 0.05). The noise ceiling, quantified as leave-one-subject-out correlation, is shown in light grey (mean ± SEM).

### EEG neural patterns correlate with responses first from early visual cortex, then TPJ/pSTS, and finally somatosensory cortex

We tracked the spatial-temporal neural dynamics of touch observation using EEG-fMRI fusion methods to examine the information flow between EVC, TPJ/pSTS, and somatosensory cortex. These results show a clear order between the neural latencies of each ROI. We find that neural information first arises in early visual cortex 50 ms post video onset, and then in TPJ/pSTS at 110 ms, and finally in somatosensory cortex at 190 ms (Figure 5, bottom, p < 0.001 for all passible comparisons). A Mann-Whitney U-test showed significant differences in all three combinations of pairs (p < 0.001).

### Visual and social perceptual, but not somatosensory, brain regions explain unique variance in EEG signals during touch observation

Given the significant correlation between neural patterns in TPJ/pSTS and those in somatosensory cortex (Figure 3), it is unclear whether the time-resolved correlations between those brain responses and EEG signals (bottom panel in Figure 5) are driven by shared or unique variance across different brain regions. Thus, we examined unique and shared contribution of fMRI responses in each region to EEG neural patterns using variance partitioning analysis to further characterize the information flow across the brain regions involved in touch observation. Variance partitioning revealed that EEG neural patterns are uniquely explained by EVC at 94 ms after video onset, then by TPJ/pSTS at 190 ms (Figure 6A). Importantly, somatosensory cortex activity did not explain any unique variance in the EEG data, but shared variance with TPJ/pSTS beginning at 206 ms (Figure 6A and B). There was also shared variance between EVC and TPJ/pSTS at about 100 ms after the initial onset of EVC, suggesting feedback from the social brain to EVC at later time points (Figure 6B).

**Figure 6.**
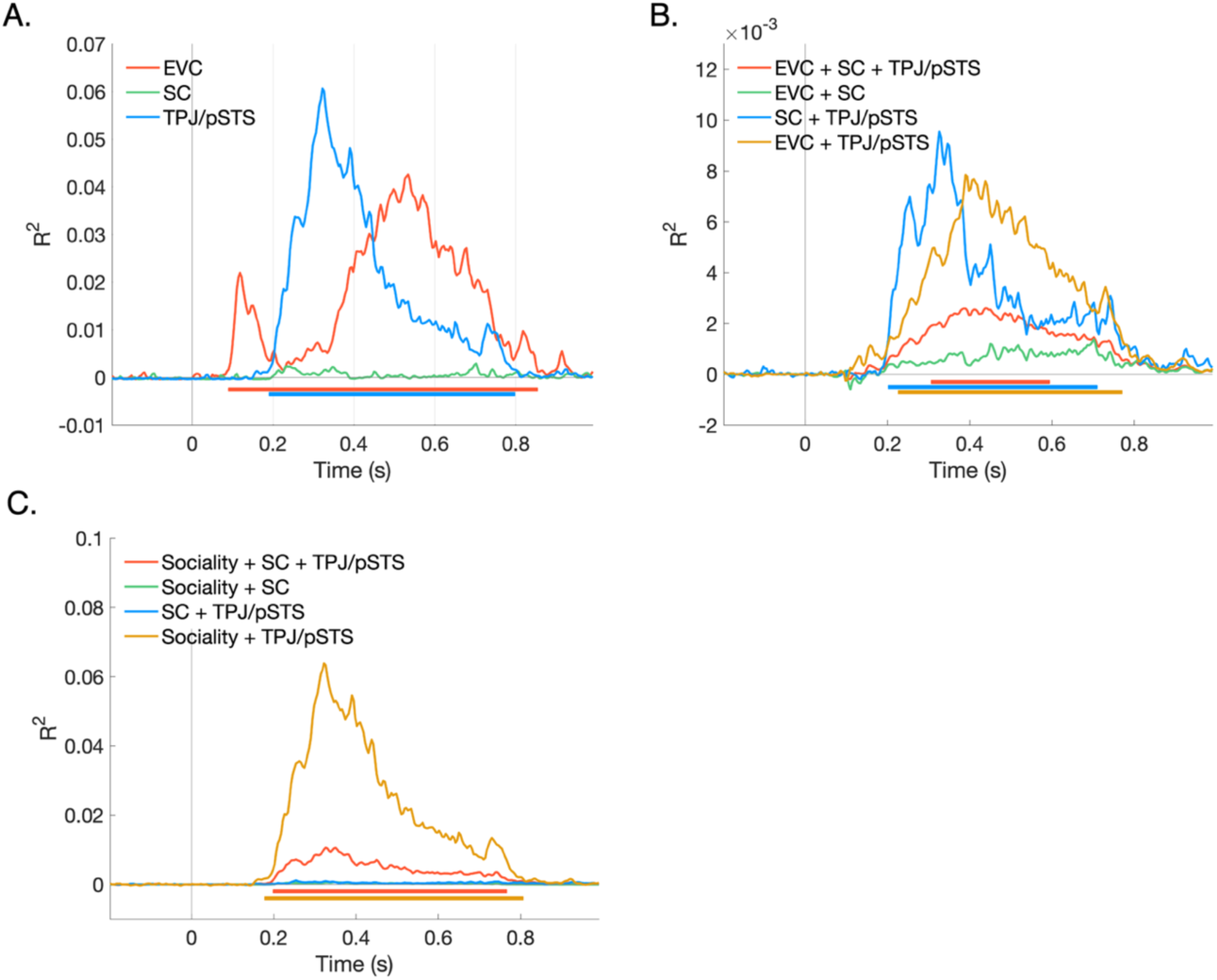
Time-resolved variance partitioning results. Multiple regressions were performed with the time-resolved EEG RDM averaged across subjects as a dependent variable. A. The unique contributions of the fMRI activity in three brain regions with EEG data. B. The shared contributions of the three brain regions with EEG data. C. The shared contributions of TPJ/pSTS and somatosensory cortex (SC) with the sociality feature. The horizontal line in each plot marks significant time windows (one-tailed sign permutation for the group-level statistics, cluster-corrected p < 0.05).

### Temporally resolved information in TPJ/pSTS explains unique variance in social touch observation

Lastly, we asked whether and when the sociality of observed touch, the feature that explains EEG data the most strongly, shares variance with TPJ/pSTS or with somatosensory cortex activity. We find that TPJ/pSTS responses share variance with sociality in explaining the EEG data at 180 ms after video onset (Figure 6C). In contrast, activity in somatosensory cortex alone does not share variance with sociality. TPJ/pSTS, somatosensory cortex, and sociality do explain shared variance with EEG signals, but this is substantially later and weaker than variance explained by TPJ/pSTS and sociality alone. Together these results suggest little direct involvement of somatosensory cortex in representing the sociality of observed touch.

## Discussion

We examined the spatiotemporal neural dynamics of social touch observation. Combining time-resolved RSA, fMRI-EEG fusion, and variance partitioning analyses (Figure 2), we for the first time identified the speed at which each feature of observed touch is processed (Figure 5), as well as the direction of information flow between brain regions. Our results revealed a direct pathway between early visual processing and social perception, with only later involvement of somatosensory simulation (Figure 6).

### Early processing of social-affective meaning of observed touch

Observing physical contact between two individuals resulted in greater neural activity in the frontal regions, while observing an individual touching an object led to stronger neural activity in the occipital regions (Figure 4). These findings are consistent with previous neuroimaging studies showing differential neural responses to social and nonsocial touch (Peled-Avron et al., 2016; Peled-Avron and Shamay-Tsoory, 2017; Lee Masson et al., 2018, 2019). Similar to previous work, the current study suggests that observing two individuals exchanging touch involves social-cognitive mechanisms, such as mentalizing and emotion recognition (Peled-Avron and Shamay-Tsoory, 2017; Lee Masson et al., 2018, 2020; Schirmer and McGlone, 2018; Arioli and Canessa, 2019), whereas recognizing a touched object requires additional visual processing (Goodale et al., 1994; Lee Masson et al., 2018, 2020; Wurm and Caramazza, 2022). Concerning the processing speed, ERPs evoked by social touch were distinguishable from those elicited by nonsocial touch as early as 100 ms after stimulus onset. Overall, the current study extends earlier findings by revealing that the neural distinction between social and nonsocial touch is established at an early stage in neural processing.

A classical ERP method is analogous to a univariate approach in fMRI, in that both methods unveil the extent to which a channel or voxel is activated in response to a given stimulus. However, this univariate method does not provide a comprehensive account of how the configuration of multiple channels evolves over time or how stimulus features are represented in the spatial patterns (Davis et al., 2014; Pillet et al., 2018). To answer this question, we employed time-resolved RSA to link EEG spatial patterns to visual and social-affective features characterizing a touch event (Figure 2). Time-resolved RSA revealed that social-affective information – whether touch is social, pleasant, or emotionally intense – is processed rapidly and immediately following the processing of low-level visual features, motion, and body movements (Figure 5).

To date, no studies have used time-resolved RSA to investigate the temporal dynamics of neural responses during touch observation. Nonetheless, our findings on the speed of visual processing during touch observation are consistent with those reported in other areas of research. We found that the low-level visual features extracted with the first layer of AlexNet explain EEG neural patterns early, beginning at 90 ms after stimulus onset. This finding aligns with previous work investigating scene perception and action observation using similar methods (Cichy et al., 2017; Dima et al., 2022). The perception of biological motion emerges early as well, beginning at 120 ms. The onset latency observed in the current study is consistent with the timing of neural processing reported in other studies of biological motion and action perception (Oram and Perrett, 1994; Chang et al., 2021; Dima et al., 2022). It is important to clarify that the term “biological motion” here does not refer to action categories. Rather, it refers to the perception of body movements. As an example, body movements required for different actions, such as hugging a person or carrying a box, are perceived similarly. Regarding motion energy, since we rely on basic optic flow techniques to calculate motion and the present study is not designed to focus on motion energy, we remain cautious in providing detailed interpretation of our results (refer to Bergen and Adelson, 1985; Dima et al., 2022).

We found that the onset latency for processing sociality, arousal, and valence information of observed touch occurred within a time frame of 150-180 ms. This finding aligns with previous research that has demonstrated the early processing of perceived valence (145 ms) and arousal (175 ms) in a variety of emotional images (Grootswagers et al., 2020). However, the processing speed of social-affective information varies and may be influenced by the level of complexity and naturalness of the stimuli used. Sociality information from simple, well-controlled images is processed rapidly, as indicated by changes in P100 components (Peled-Avron and Shamay-Tsoory, 2017), whereas natural stimuli tend to require more time to process (Isik et al., 2020; Dima et al., 2022). The stimulus set used in this study focused on body movement without facial information and other contextual information about touch, which may explain the fast processing times. Further research is needed to investigate the speed at which social-affective information is extracted in more ecologically valid settings, where contextual details about touch may play a critical role in comprehending its meaning. Taken together, the ERP and RSA techniques showed that the brain detects the social-affective significance of touch at an early stage, well within the timeframe of feedforward visual processing (Lamme and Roelfsema, 2000). These results highlight the importance of touch perception as a fundamental aspect of the human visual experience.

### The social-affective meaning of touch is initially extracted through a social perceptual pathway

With time-resolved RSA and variance partitioning analyses, we determined the direction of information flow between EVC, TPJ/pSTS and somatosensory cortex. We demonstrated that the social-affective meaning of touch is initially extracted through a social perceptual pathway, followed by the subsequent involvement of somatosensory simulation. These findings suggest that touch observation is not mediated by the reenactment of somatosensory representations, which contradicts the embodiment theory of social perception (Rizzolatti and Sinigaglia, 2010, 2016). Instead, the somatosensory cortex appears to receive information from a social perceptual pathway, as indicated by this region’s absence of unique variance with EEG neural activity.

Although somatosensory involvement occurs at a later stage, its contribution to social understanding should not be undervalued. Individuals who struggle with social interaction, particularly in the context of interpersonal touch, tend to exhibit weak or absent somatosensory activity, which can decrease social bonding (Peled-Avron and Shamay-Tsoory, 2017; Lee Masson et al., 2018, 2019). The relay of social information from TPJ/pSTS to the somatosensory cortex may help individuals empathize with others and comprehend another person’s emotional states at a more profound level (Schaefer et al., 2012, 2013; Bolognini et al., 2013; Peled-Avron et al., 2016; Peled-Avron and Woolley, 2022). The methodology presented here provides an opportunity for future work to examine whether the lack of somatosensory simulation in particular groups of individuals is associated with inadequate information flow between social perceptual and somatosensory system.

## Conclusion

Our study, for the first time, revealed that social-affective information of observed touch is processed rapidly and directly through social perceptual brain regions. Social touch plays an important role in establishing and maintaining social bonds (Hertenstein et al., 2006b; Chatel-Goldman et al., 2014; Suvilehto et al., 2015). It facilitates effective communication, fosters greater trust and empathy, and provides relief from stress, anxiety, and pain, ultimately leading to improved psychological and physical wellbeing (Debrot et al., 2013; Korisky et al., 2020; Shamay-Tsoory and Eisenberger, 2021). Our findings suggest that the early neural processing of social and emotional signals conveyed through touch may play a key role in the successful use of social touch during rapidly-changing, real world social interactions.

## Acknowledgements

The authors wish to thank Lucy Chang, Victoria Liu, and Diana Dima for their help with the EEG data collection, Diana Dima for sharing her code for EEG analysis, and Diana Dima and Emalie McMahon for feedback on the manuscript. This work was supported with funds from The Clare Boothe Luce Program for Women in STEM.

